# Direct morpho-chemical characterization of elusive plant residues from Aurignacian Pontic Steppe ground stones

**DOI:** 10.1101/2020.07.23.212324

**Authors:** G. Birarda, C. Cagnato, I. Pantyukhina, C. Stani, N. Cefarin, G. Sorrentino, E. Badetti, A. Marcomini, C. Lubritto, G. Khlopachev, S. Covalenco, T. Obada, N. Skakun, L. Vaccari, L. Longo

## Abstract

Direct evidence for the intentional processing of starch-rich plants during the Paleolithic is scant, and that evidence is often compromised by concerns over preservation and contamination. Our integrated, multimodal approach couples wear-trace analysis with chemical imaging methods to identify the presence of genuine ancient starch candidates (ASC) on ground stones used in the Pontic Steppe starting around 40,000 years ago. Optical and electron microscopy coupled with infrared spectromicroscopy and imaging provide morphological and chemical profiles for ASCs, that partially match the vibrational polysaccharide features of modern reference starches, highlighting diagenetic differences ranging from partial oxidation to mineralization. The results suggest the intentional processing of roots and tubers by means of mechanical tenderization and shed light on the role of dietary carbohydrates during Homo sapiens’ (HS) colonization of Eurasia, demonstrating a long acquaintance with predictable calorific foods, crucial to maintain homeostasis during the harsh conditions of the Late MIS 3 (40-25 ky).

## Introduction

Starch is how plants store energy, and a highly energetic and nutritious food for humans. The consumption of starch-rich storage organs has been documented since the Middle Pleistocene through the extraction of starch grains from dental calculus, coprolites, and gut contents, which can be considered as direct evidence of their role in the diet ^1–4^. On the other hand, charred roots and tubers recognized at early modern human sites in South Africa and in northwestern Australia^5–7^, and starch grains retrieved in sediments from Klissoura cave in Greece ^8^ represent indirect proof of starchy plant foraging. The Epipalaeolithic site of Ohalo II on the Galilee lake yielded a unique record of thousands of charred protoweeds and other plants remains as well as ground stones used to process starchy plants ^9,10^.

In order to enhance the nutritional properties of starchy plants they must be physically processed by grounding and pounding, and eventually thermally treated to release their nutritional bioavailability to, finally, generate energy. Ground stones are direct evidence for human-induced mechanical tenderization of starchy organs. Indeed, starch grains were recognized on a handful of flint flakes from Layer Fa of Payre (Rhone Valley, France, beginning of MIS 3) and on one trapezoid flint tool from the Early Upper Palaeolithic Layer C of Buran Kaya III (Crimea)^11,12^. More consistent evidence of intentional starch processing emerged during the Gravettian, when grinding stones and pestles from the Italian peninsula and central Europe sites clearly were used to mechanically tenderize underground storage organs (USOs) to obtain a coarsely-ground flour ^13,14^. Charred plant remains used as food are also reported for late Mesolithic sites from northern Europe (Ertebolle, a coastal settlement in southern Scandinavia ^15^ and Lubuskie Lakeland in Poland ^16^).

However, little is known about the intentional processing of starchy plants during the earliest colonization by HS of a totally new environment - the northern Eurasian continent - under the very challenging climatic downturn due to the “volcanic winters” between ca 50 to 38 ka ^17,18^. For the present study, we selected two southern Pontic steppe sites - Surein I (a rockshelter in Crimea) and Brinzeni I (a cave in Moldova) - driven by the following considerations: their chrono-cultural attribution to the Aurignacian, listed among the oldest evidence of HS occupation in the area perhaps as a result of this area being a *refugium*, and the richness in ground stones and the associated presence of human teeth attributed to HS ^19,20^. Brinzeni I counts more than 100 among flat slabs and pebbles putatively interpreted as ground stones retrieved in Layer 3 with related excavation details (mapping, provenience), however we focused our attention on a selection of 36 percussive tools and here we present the in-deep analysis of 8 among grinding stones and pestles. The large slab from Surein I was associated with a structure in square 9B/4-5 from Layer 3, photographed and mapped during the Bonch-Osmolovsky 1926-29 excavation ^21^ and philologically displayed at MAE RAS (St. Petersburg). The ground stones under inspection are part of a broad and innovative research design devoted to EUP percussive tools, possibly used to process dietary carbohydrates, described by the authors ^13,14,22–25^.

The ground stones, excavated at two Aurignacian sites in the Pontic Forest-Steppe were examined from the functional and use-related biogenic residues perspective by coupling wear-traces and associated starch grains analyses, anticipated by a thorough cleaning of the used surfaces with a multi peeling process, to avoid putative contaminants (see Methods section). The research design relies in the observations of the ground stones from the macro-scale (3D scans and photogrammetry), to optical and digital microscopy, down to the nano-scale by means of scanning electron microscopy (SEM) ^23,24^, and integrates the physical-chemical characterization of the associated biogenic residues. Starch granules recovered from these ground stones were investigated by: (i) light (Optical) and (ii) scanning electron microscopies, as well as by (iii) FTIR spectromicroscopy and imaging with both conventional infrared source and (iv) infrared synchrotron radiation (IRSR), for providing high resolution chemical profiles of a single starch grain. This is the very first time that these aforementioned techniques are applied to study Palaeolithic starches.

Our results demonstrate the synergistic co-occurrence of the intentional processing of starchy plants with the earliest settling in western Eurasia by modern humans. The interdisciplinary methodology applied to the observed ASCs provides experimental evidences for interpreting the Use-Related Biogenic Residues (U-RBRs) from Brinzeni I and Surein I as indeed composed of ancient starch ^26,27^ and attribute their consumption to modern human dwellers approximately 40.000 years ago. Our various lines of data that include the co-presence of U-RBRs (starch, phytoliths, fibres, raphides), on Aurignacian ground stones is bolstered by FTIR imaging and multivariate spectral analysis. Our multidimensional contribution establishes an investigative procedure that combines morphological and chemical analyses, recognizes and authenticates ancient use-related starch granules, and sheds light on the origins of starchy food processing in south-eastern Europe during the Aurignacian.

### Archaeological context of the research

The Pontic steppe covers the western part of an immense space (the Eurasia Steppe Belt) which crossed east-west Eurasia, in a mosaic of river valleys from the Dnepr to the Volga and to the Anui, and high plateaus from the Carpathians to the Urals to the Altai Mountains. Its southern rim covers the Mediterranean coastal areas of the Euxeinos Pontos, as it was commonly known in antiquity, overlooking the Black Sea and the Caspian Sea and functioning as an open nexus between the northern and southern boreal territories. This biome is a rich steppe-like grassland dominated by shrubs with spots of forest-steppe. Regarding the carrying capacity of the territory entered by HS around 40,000 years ago ^28^, the biotic diversity is manifested across the agencies that populated the Pontic Steppe. Different taxa of grazers and browsers (mega and large herbivores, mammoth, woolly rhinoceros, bison, horse, wild ass, saiga, deer including giant deer, all source of fats and proteins for the carnivores) were traditional suppliers of fats and proteins. This wide faunal spectrum is indicative of a varied geomorphology that included river valleys and steep slopes where the caves opened up, and in turn reflects the plants on which they fed. It is our point that small mobile groups of Late Pleistocene hunter-gatherers strategically foraged on a broad spectrum of resources that included starchy plants. The European south-eastern territories, such as Moldova and Crimea, were peripheral to the permafrost and variations in sea levels allowed for east-west migrations towards patchy forested landscapes ^29^. In these terms, southeastern Europe would appear as a *refugium* for both humans and animals, as evidenced by the richness of late MIS 3 (40-25 ky calBP) settlements occupied during the latest phases of the Middle Palaeolithic (MP, Micoquian) and the Early Upper Palaeolithic (EUP, Aurignacian, ^30,31^. Both the Prut River territory and southern Crimean outcrops were rich in water sources, caves and rock shelters, raw material to be transformed into tools, and in both animal and vegetal foods. Here we report on percussive stone tools used to mechanically process starchy plants. Our hypothesis is that small mobile groups of Late Pleistocene hunter-gatherers strategically foraged on a broad spectrum of resources that included starchy plants.

The nine ground stones investigated in this study were retrieved at Brinzeni I, a cave on the Prut river basin (Moldova) and at Surein I, a rock shelter on the southern slopes of Crimea ^23,24^. As shown in Figure 1 A-B-C, the sites are located within a crucial territory that set the scene for the early occurrence of HS in the *refugia* areas of the northern rim of the Mediterranean Sea, who most probably crossed paths and mated with local late Neandertals as supported by the remains at Oase cave (Romania) and Bacho Kiro (Bulgaria) ^28,32^. In spite of cross-breeding, paleogenomics currently supports a marked difference in terms of starch food bio accessibility among the two hominins, with a clear positive selection of gene clusters advocating for an increased efficient metabolization of dietary carbohydrates in HS ^33,34^. Therefore, south-eastern Europe and the Crimean peninsula became one of the key regions to study the dietary strategies during the coexistence of Neandertals and modern humans ^35,36^.

**Fig. 1.**
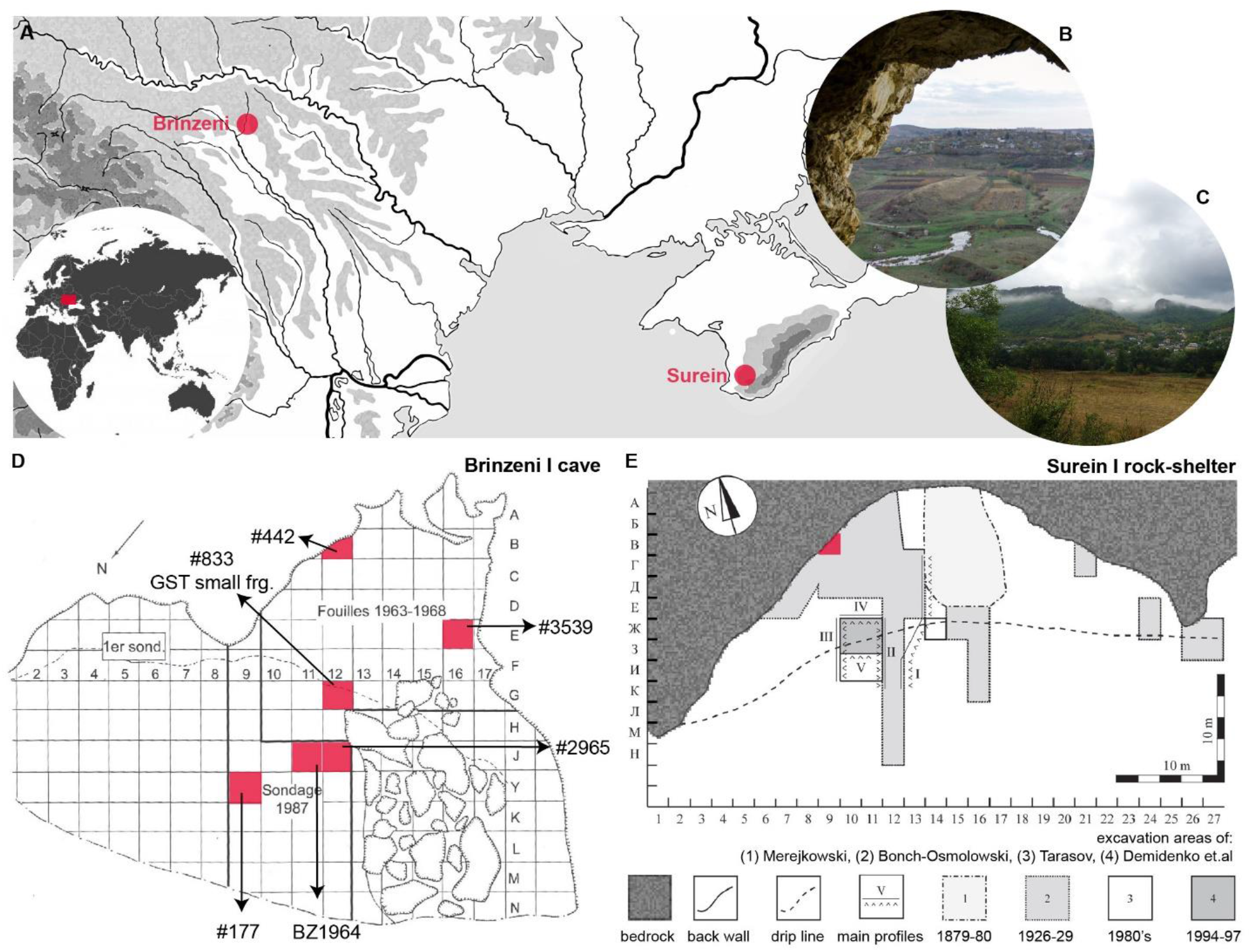
Geographic localization of the selected sites. **(A)** Pontic Steppe where the sites are located. **(B)** view from the inside of Brinzeni I cave overlooking the Rakozev River valley, a tributary creek of the Prut river; **(C)** View of the Bel’bek River gorge on which the Surein cave opens; **(D)** Brinzeni I site planimetry where the investigated ground stones are mapped (modified from Allsworth-Jones et al. 2018, but maintained the Cyrillic alphabet). Red squares yielded the investigated ground stones: #442 in square 12 B, #no number GST small fragment in square 12 Γ, #833 in square 12 Γ, #2965 in square 12 Ж (these four fragments refit into a grinding stone and a complete pestle); #No number BZ1964 in square 11 Ж, #177 in square 9 И; #6707 (no square provenience) is a large silty sandstone, #3539 in square 16 E, is a small limestone slab. **(E)** Surein I planimetry (Bonch-Osmolovsky excavation 1926-29, modified from Vekilova, 1957), the limestone slab was retrieved in square 9 B, layer 3.

## Results

### Ground stones analysis

The grinding stone from Surein I is a large oval slab made of biogenic limestone retrieved in the lowermost layer 3 square 9 B (Figure 1E), and has an active surface with wear-traces and associated starch grains ^23^. From the large assemblage of Brinzeni I, five grinding stones and three pestles have been analysed (Figure 1 D). Of these, one grinding stone (#442 square 12 B #no number square 12 Γ,) and one pestle (#833 square 12 Γ, #2965 square 12 Ж) were broken and the refitting was made during this study. The remaining ground stones are mostly made out of silty sandstone with small size quartz grains (#177 square 9 И; #no number US 5 square 11 Ж); #6707 (Layer 3, but exact square of provenience unknown) is a large silty sandstone and #3539 (square 16 E) is a small limestone slab ^24^. Differently from Holocene ground stones, the percussive tools used during the EUP are unshaped pebbles, although a certain standardization is evident in the selection of size, shape, and raw materials. Wear-traces were recognized on the active areas where the contact with the working materials was more prolonged or intense, affecting the salient parts of the uneven surfaces, in the form of spotted polish, alignment of striations, and isolated striae (Figure 2). Functional analysis was carried out coupling optical and digital microscopy^22^. We established a specific procedure to extract use-related starch grains from these areas with the purpose of coupling physical-chemical characterization of starch grains, previously un-attempted (see Methods below, ^23,37^).

**Fig. 2.**
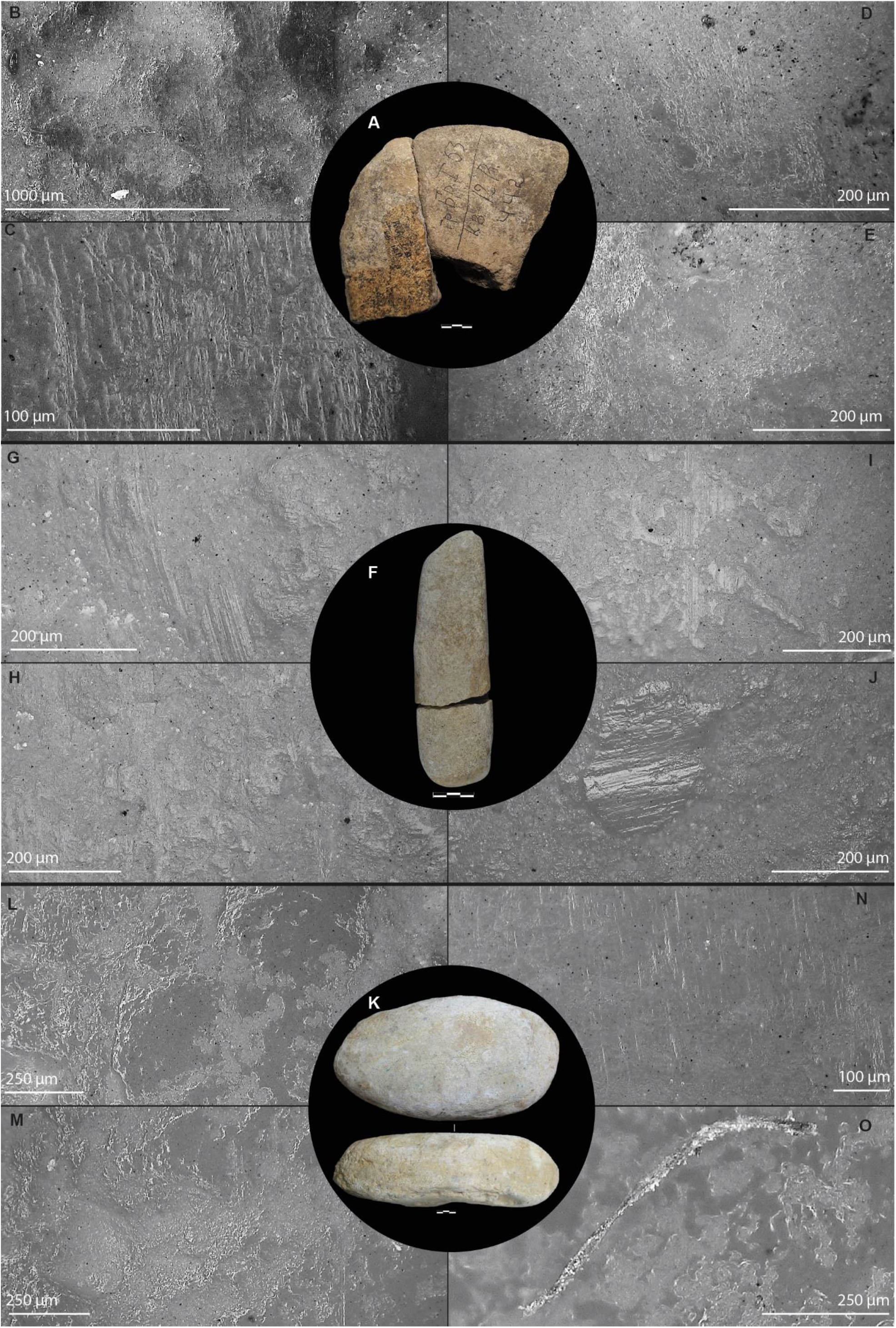
The Ground stones and the wear traces analysis. **(A-J)** Brinzeni I: **(A)** wear-traces on the refitted grinding stone and the pestle, **(B-C)** #No Number, left fragment, **(D-E)** #442, right fragment, **(F)** Broken pestle, **(G-H)** #833, upper fragment, **(I-J)** #2965, lower fragment. In panels **(B-C-D-E-G-H-I-J)** Aligned polished areas and striations. (**K-O)** Surein I grinding stone, Face A (several used areas) and lateral view; **(L-N)** spotted polish, on which linear traces and alignment are visible fashioned as shallow lines of varying lengths. **(O)** Polished areas associated with a fibre. Digital microscope Hirox KH-8700 (lens MXG-2500REZ).

The wear-traces analysis of the mechanical processing of starchy plants by means of ground stones was first carried out directly on the stones in museum collections, and then by using imprints to reproduce the stones’ surface texture at the nanometric scale further continued off-site, in laboratory settings (see below: Methods). This procedure, which included 3D scans, Optical, Digital (DM) and SEM microscopy aimed at resolving the function(s) of the percussive tools under scrutiny and disclosed details and served as a complementary source of surface analysis for both artefacts and starches. The observation of the molds using DM and SEM revealed the contextual recognition of plant residues and specifically starch grains still adhering to the used areas and can be considered a step-change methodological refinement in U-RBR analysis.

Functional analysis detects the presence of several wear-traces (detailed elsewhere, ^23,24^) which we briefly recall here and present in Figure 2. The raw materials of the two assemblages are different: a micritic sandstone rich in micrometric quartz crystals characterizes the grinding stones and the pestles (hand-held active tools) from Brinzeni I cave (Moldova) while the Surein I (Crimea) tool is a large steady biogenic limestone grinding stone. The two raw materials influence the degree of development of wear-traces and how they are featured: weak glossy spots and spotted polish with varying degrees of brightness, which, under the DM appear matte and darkish grey as the result of a flattening of the surface’s uneven micro relief (Figure 2 E, L-M, O); clearly aligned parallel linear traces that cross the image (Figure 2 B-D; I, J, N). Other traces include more diffused spotty polish associated with striations that affect the salient parts of the micro topography (Figure 2 G-H; L-M) and the presence of entrapped fibers on the striated areas on the most prominent reliefs (Figure 2 O).

The Brinzeni I pestles showed more intense traces on the rounded apex of the movable tool which was used for pounding and threshing activities usually in bi-directional coupled kinematics. The large broken grinding stone from Brinzeni I shows the highest development of wear-traces in the sections where the two fragments refit, clearly demonstrating this is the part undergoing maximum mechanical stress. For the Surein I grinding stones, the most-defined use-wear traces are concentrated in the central part and, to a lesser extent, in peripheral areas.

### Use-related starch grains

We performed the high-resolution physical-chemical characterization of ancient starch grains after their extraction from actively used areas of the ground stones according to the authors’ standard procedure (Methods below, ^23,24^). In detail the analysis focused on the extracted material that present at least two of the following characteristics: i-starch-like morphology as revealed by SEM and/or optical analyses, ii-Maltese cross when inspected with polarized light, iii-main FTIR spectral features of polysaccharides ^38^. These extracted particles will be referred to as “ancient starch candidates”, ASC, to distinguish them from Modern Starch Reference (MSR). A finer classification of both ASC and MSR is reported hereafter:

- **ASC-1**: Starch observed directly on the molds. Not suitable for FTIR analyses.
- **ASC-2**: Encrustations adhering to the ground stones removed by sonication of the stone and crushed using a laboratory agate mortar and pestle until obtaining a fine powder that was then suspended in ultrapure water and deposited on a ZnSe window for FTIR measurements.
- **ASC-3**: Starch isolated from the sonication of molds in ultrapure water. The particle suspension was deposited on a silicon or ZnSe window for FTIR measurements.
- **ASC-4**: Starch isolated according to published protocols (Pearsall et al. 2004).
- **MSR-1**: Starch grains extracted from grinding and pounding reproducing ancient technologies.
- **MSR-2**: Laboratory processed starch grains obtained by sequential water and ethanol extractions.

Optical microscopy (Figure 3) and SEM data (Figure 4) will be presented in order to provide insights on the morphology and fine structure of the recovered ASCs, and of other plant remains observed in the samples. Then, FTIR data on dried powder deposits (ASC-2 and ACS-3) and isolated ASC-4 will be commented in order to delineate the chemical profile of the particles of interest. Then the ASCs FTIR spectra will be compared with those of MSR-1 and MSR-2, and confronted using a chemometric model.

**Fig. 3.**
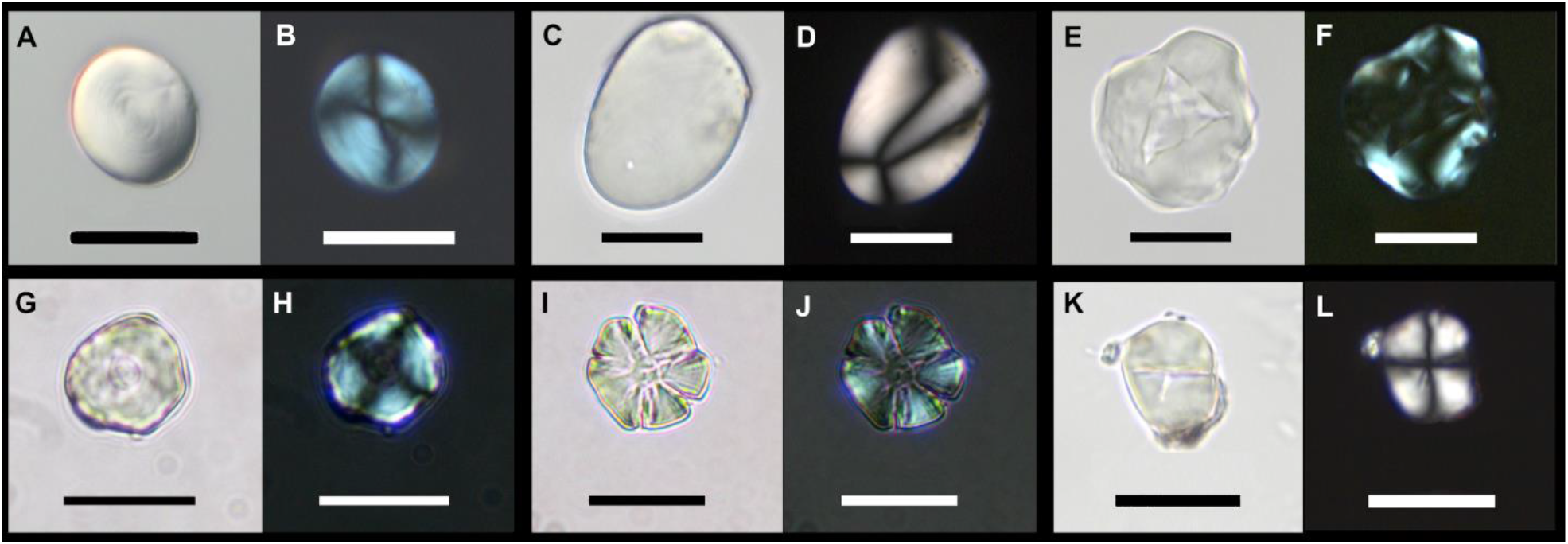
Optical characterization of the ASCs. **(A-H)** Starch grains from Brinzeni I and **(K-L)** Surein I under Optical Microscope direct and polarized light. Starches from Brinzeni: **(A-B)** Sample 5, **(C-D)** #177, **(E-F)** #833, **(G-H)** #3539, **(I-J)** #6706, and from Surein I **(K-L)**. Scale bars 20 microns.

**Fig. 4.**
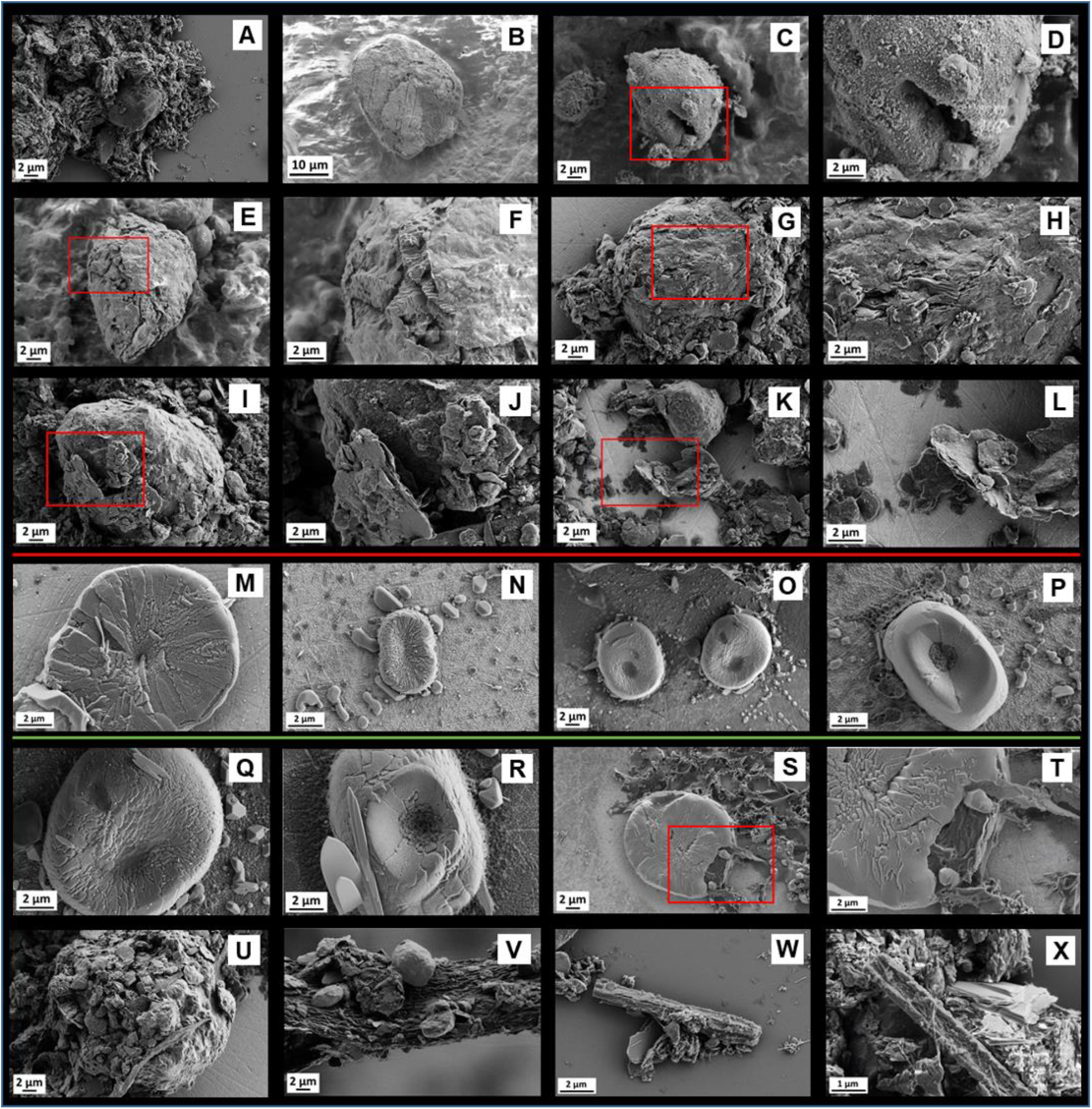
SEM characterization of the ASCs and U-RBRs. SEM micrographs of starch grains extracted from different ground stones. **(A-L)** Brinzeni I, and **(M-P)** Surein I separated by a red line. The green line separates the panels dedicated to the morphological characterization of starches and their modifications, from the Use-Related Biogenic Residues (U-RBR) belonging to plants. **(Q-R)** Raphides directly associated with starches are evident in samples from Surein I, in **(S-T)** parenchyma remains are shown, in **(U-V)** some fibres, with some starch grains attached (see also S1) and in **(W-X)** phytoliths.

### Use-Related Biogenic residues: Optical and SEM characterization

Starches are synthesized in amyloplasts and deposited in rings composed of amylose and amylopectin that grow as grains ^39,40^. Grains are then stored in various organs of the plant, both in underground storage organs (USO) such as roots, rhizomes, and tubers, and above ground organs (ASO) that include fruits and seeds.

Through the optical microscopy inspection, several use-related plant remains, namely starch grains (Figure 3 and Figure 4) and fiber (Figure S1), were identified on both the ground stones from Brinzeni I and Surein I, according to standardized procedures as reported in ^24^. Then, use-related residues were observed directly on the molds by means of Digital (Figure 2 O) and SEM microscopy (ASC-1, Fig. 4 B-C, Figure S1 E-F). Afterwards, some particles were separated from the stones by soaking the targeted used areas of the tool in an ultrasonic tank cleaner at room temperature (ASC-2). Intriguingly enough, the SEM preliminary inspection revealed that U-RBR namely starch grains and fibers were still adhering to the imprints. Once U-RBR were observed still adhering to the molds, a second approach used the molds as a source of plant residues and even the molds (or selected parts of them) were sonicated to extract starch grains (ASC-3). The procedure to extract ASC-4 has already been described in detail elsewhere ^23,37^.

Overall, from the samples whereby the starch grains were isolated using standardized laboratory protocols, e.g. ^41^, and using optical microscopy, we counted a total of 66 starch grains on the tools from both Brinzeni (n=59) and Surein (n=7). The size of the grains is micrometric, averaging less than 50 μm in the case of our samples. Light microscope resolution, up to 0.2 μm, allows for viewing the morphological features as well as the characteristic extinction cross (Maltese cross) evident under cross-polarized light. Some examples of recovered ASC-4 starches are shown in Figure 3. We were unable to determine their taxonomic identifications, due to the relative poor conservation of a large majority of the starch grains, but also because of a lack of reference collection for this geographic region and time period ^12,42^. However, it was possible to distinguish different morphologies and to detail diagnostic features namely the hila and lamellae. Moreover, several starches were clearly broken as the result of mechanical processing, and show pits and cavities interpreted as the result of biochemical activities (soil enzymes or other biogenic agents like fungi or bacteria). From Brinzeni I, we found a wide variety of morphologies, which include lenticular (Figure 3 A-B), polyhedral (Figure 3 E-F), and roundish starch grains (Figure 3 G-H), but this could be related to the larger number of samples from Brinzeni cave compared to Surein. We also recovered starch grains of probable USOs (Figure 3 C-D). From Surein, we observed polyhedral grains, which range in size between 15 and 23 μm, as well as more oval forms, which measured between 18 and 19 μm wide (Figure 3 K-L). Damages were evident on a large majority of the starch grains and include broken or crushed grains (Figure 3 I-J), deformed grains, loss of extinction crosses, as well as circular or uneven depressions affecting the central parts of the grains where the hilum is located (Figure 3 E-F). These damages suggest different types of mechanical forces or processing activities, or even taphonomic processes ^43–46^.

ASCs were also observed by SEM, revealing finer details regarding the shape, size, and overall surface details (Figure 4). Moreover, SEM allowed for the smallest sized grains to be spotted (<5 μm) which is challenging with optical microscopy. The SEM analyses were first carried out directly on the molds from both sites (ASC-1) and on the resulting dispersion of particles from the sonicated stones and molds, deposited on Si or ZnSe windows (ASC-2 and ASC-3). A large number of ASC-2 from Brinzeni was identified (Figure 4 A to L). Although they are often surrounded by an important quantity of sediment, as shown in panel A, they are clearly recognisable by their polyhedral or rounded shape. Their surfaces are not as smooth as in the case of the modern starches (see Figure 5 panel Ji). The archaeological starches exhibit rough surfaces possibly due to different kinds of damages and aging as well. The rounded starch grain shown in panel B displays some striations; these could be interpreted as the result of mechanical pounding hinting to an intentional processing of starch-rich storage organs. The starch grain in panel C shows an eroded and pitted surface. Moreover, the characteristic hilum can be appreciated, even in lateral positions (Figure 4 C-D). Some grains from Brinzeni showed cracking and exfoliation of the external surface and the typical internal amylose-amylopectin lamellar structures are exposed (Figure 4E and its magnification, panel F; panel G and its magnification, panel H). Several grains showed exfoliations and disruptions (Figure 4I and its magnification, panel J and panel K and its magnification, panel L). Grains from Surein I (Figure 4 M to T) are ASC-3 type. They appear to have less soil residues and show a smoother surface if compared with the Brinzeni ones. They are mainly characterized by an oval or roundish shape and in some cases they showed a radial fibrillar fracture surface (Figure 4 M and N), typical of crushed starch granules ^47^. A peculiar conical crater has also been observed in the major part of the Surein I grains (Figure 4 O, P and Q, R).

**Fig. 5.**
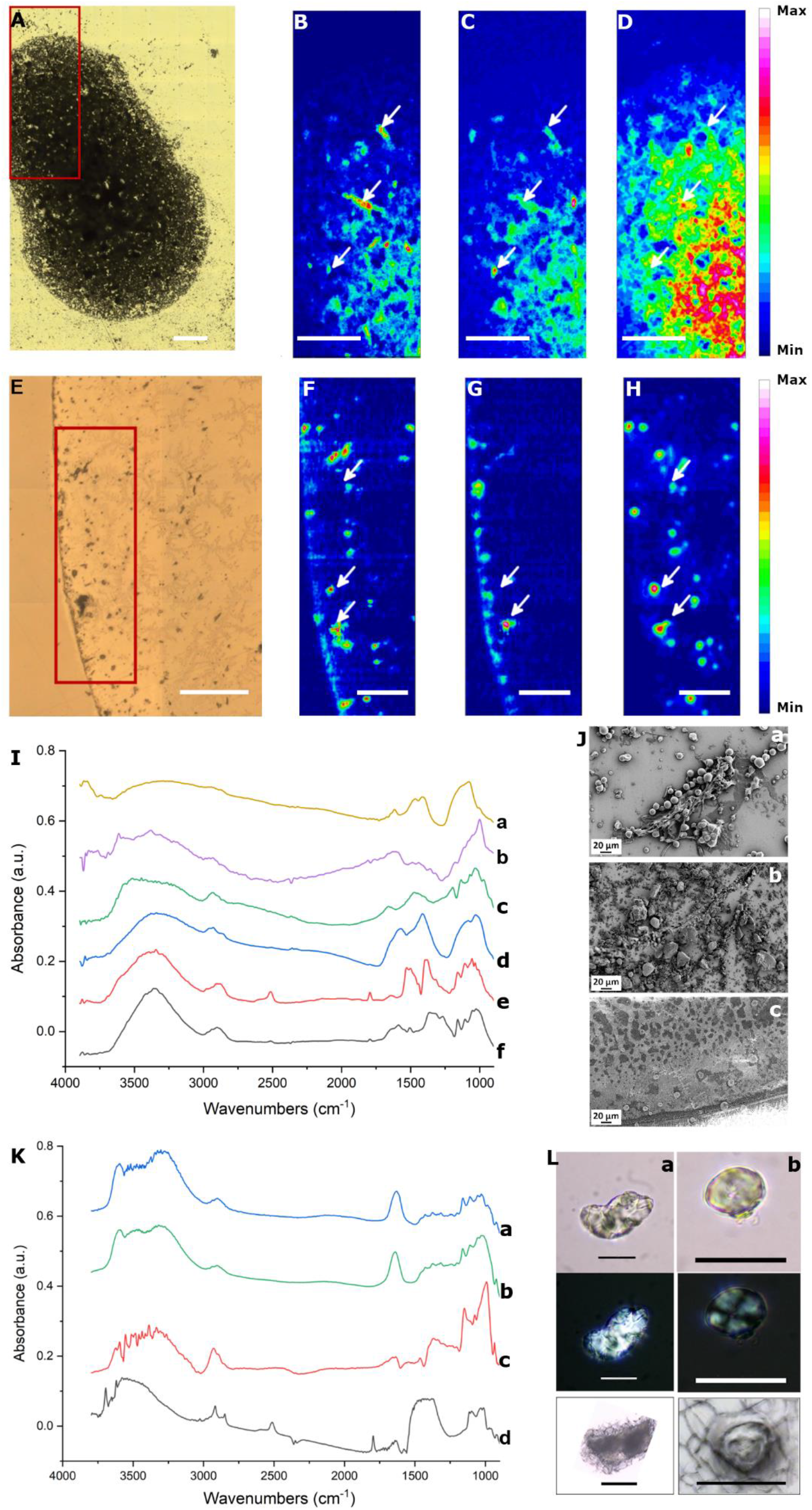
FTIR data on the ASCs. **(A)** Overview of a deposit of particles obtained from a ground stone from BZ 1964 Layer 3 square 11j (no number, Sample 5), the red rectangle indicates the measured area, the scale bar is 150 microns. **(B-D)**. Heat maps of the inspected area from Brinzeni #833 were generated by the integration of specific infrared bands: **(B)** −OH stretching, **(C)** C-H stretching, **(D)** C-O-C stretching. The scale bars are 150 microns. **(E)** overview of a drop of the suspension obtained by the sonication of mold n. 3 from Surein I, the red rectangle identifies the measured zone. The scale bar is 150 microns **(F-H)**. Heat maps generated from Surein I data as done for the Brinzeni sample. The hot spots (red-purple) represent the pixels where the integral value is higher, dark blue areas correspond always to 0. the scale bars are 75 microns. **(I)** Six representative spectra of ASC-2 and ASC-3 **a)** to **e)** from Brinzeni #833 and **f)** from Surein I. **(J)** SEM micrographs of ASCs: **a)** image of modern starches from a fresh *Manihot* root from Tanzania, **b)** image of starches surrounding a fiber from Brinzeni I, **c)** image of a group of starches from Surein I. **(K)** Spectra of some of the ASC-4: **a)** and **b)** from BZ#3539, **c)** from BZ#6707 and **d)** from BZ#442. **(L)** From top to bottom, transmitted and polarized light photographs of the starches **a)** from BZ#2965 and **b)** from BZ#6707 on the glass slide, in the lower panel, the same starches after being transferred onto the ZnSe window.

SEM images also revealed the presence of numerous U-RBR belonging to plants (Figure 4, panels Q to T from Surein I and U to X, from Brinzeni). Among them raphides, or needle-shaped calcium oxalate crystals (panels Q and R), are common in higher and lower vascular plants and algae ^48^. Based on the two symmetrical pointed ends of the crystals, it would appear that these belong to Type I, considered the most common form of raphides ^49^. However, to confirm this we would also have to see them in cross-section and verify they are indeed four-sided, which we were unable to do. While it is not possible to identify these raphides to their taxon, their presence further supports our argument that the ground stone tool was used to process plant matter, arguments put forward by Hardy and colleagues ^12^ following their research at the site of Buran Kaya III in Crimea.

Residues of amyloplast parenchyma are dispersed in several samples both from Surein I (Figure 4, panel S, surrounding the starch granule; panel T) and Brinzeni (panel U, light green colored). We also noticed that the starch granule in panel S, magnified in panel T, showed a different pattern of degradation if compared with starches from Brinzeni I. If the latter presented damages to the external surface, this starch from Surein I revealed to be affected by erosion or solubilization of the internal portion whilst keeping the external surface almost intact, even if crushed and flattened as already observed for the granules from this site (see details in panel T).

We also noted the presence of a fiber on the Brinzeni I grinding stone (small fragment), which seems very withered and covered in both minerals and a couple of starch grains (Figure 4V, and supplementary materials). Finally, putative elongated phytoliths (silica bodies) produced within both the intra- and extracellular structures of plants, were observed, especially in the samples from Surein I and from Brinzeni I, sample 5 (pestle) (Figure 4, panels W and X).

Furthermore, with the level of detail reachable with this imaging technique, it was possible to see the fine structure of the ASCs down to the individual lamellae that constitute the starches (Figure 4F and H). Nevertheless, to reach beyond the descriptive approaches up to now employed in the analysis of this kind of samples, we complemented the morphological analyses with a label free, non-damaging chemical characterization of the samples by using FTIR microscopy and imaging.

### FTIR imaging on particles from the ground stones

Several far field IR images were collected and then the data processed and analysed. As can be seen from optical images in Figure 5A and 5E, showing ASC-2 sample obtained from the Brinzeni pestle BZ#833 and ASC-3 sample from Surein I, the sonicated powders present a predominance of mineral signals, with few organic materials mixed with soil particles. Several round, starch-like objects can be observed, although not all of them have a FTIR spectrum, that is a chemical composition, compatible with a carbohydrate-based particle ^50^.

As a matter of fact, aiming to find and characterize any ASCs, chemicals images were generated by integrating the infrared hyperspectral data at specific spectral intervals indicative of carbohydrates: 3500-3100 cm^−1^ for the OH stretching, 3000-2800 cm^−1^ for −CH stretching of methyl and methylene groups, 1200–900 cm^−1^ for the C-O, C-C and C-OH stretching of carbohydrate backbone^51^. Conventionally, this latter spectral range is the most distinctive for carbohydrates, and used for their characterization. Nevertheless, at the same energies there can be an interference due to the presence of signals from metal-oxides, silicates, phosphates and sulphates from the soil that, indeed, generate “false” hotspots in the chemical maps ^52^.

By observing the chemical images generated integrating the 1100-1150 cm^−1^ spectral region in Fig. 5D and H, it is clear how they portray among the all possible ASC hotspots, but also surely the minerals from the soil. Especially, in the deposition from Brinzeni, the particles are so densely packed together, making it really difficult to identify the single starches in the C-O-C chemical image (Figure 5D). Nevertheless, by comparing the same image pixels in the other two chemical maps (B and C), it is possible to identify areas with common local maxima, pointed by arrows, where also the other peculiar carbohydrate signals are intense. From these areas, average spectra of ASCs were extracted and some of them are presented in panel 5I. It has to be highlighted at this stage that, in order to bring out the ASC chemical profile over the soil contribution, the spectral contribution of the surrounding material has been carefully subtracted, as described in methods. The spectral profiles of these ASCs present strong absorbance signals in the 1200900 cm^−1^ range, typical of C-O-C, C-O and C-C ring vibrations, medium to strong −OH signals, index of a different degree of degradation/oxidation, and weak to medium −CH stretching signals, index of a partial chain scission, with the main peak of the methylene moieties centered between 2920 and 2925 cm^−1^ for all samples. It can be noticed that some ASCs spectra, e.g. red and black spectra in Figure 5I, present signals from calcium carbonate, like the broad intense band at ~1430 cm^−1^, the C=O stretching at 1790 cm^−1^ and the overtone at 2520 cm^−1^, while some clay-like signals at higher frequencies, with sharp peaks at 3612-3614 cm^−1^, can be seen in the green and violet spectra in Figure 5I. Spectra e and f in panel 5I present also a shoulder at ~1322 cm-1, assignable to calcium oxalate. Even after subtracting the contribution of the surrounding material, it was not possible to remove these spectral features, thus it is possible to hypothesize that these signals do not come from a contamination from the soil, but are due to a partial mineralization of the ASCs due to a slow exchange with the stone of the pestle, as confirmed by ASC-4 analysis (see later/below page when definitive).

Summarizing, among all twenty-five ASCs identified among ASC-2 and ASC-3 samples inspecting over fifty chemical maps, common ASC carbohydrate-distinctive traits were identified but also a quite high variability in the peak positions of the signals in the 1100-900 cm^−1^ region as well as in the relative intensities of the main bands was observed, possibly due either to different degrees of order/crystallinity/aging or to the different plant species from which the ASCs originated from ^51,53^. As an additional note, the small number of found ASCs should suggest the absence of contamination from modern starches.

Moreover, in some samples it was possible to chemically identify other plant materials, such as small wooden fibers for example, e.g. spectrum f presents also a signal at 1511cm^−1^ from the aromatic moieties of lignin and ferulic acid. SEM and FTIR images of a fiber are shown in the supplementary materials. The fiber looks withered and covered in minerals, nevertheless the FTIR signals are preserved. Analyzing the SEM micrographs made it also possible to notice a starch particle attached to this fiber. The good preservation of the fibers retrieved on both sites ground stones strengthens our claim regarding the good preservation of other plant material in these samples (Figure S1).

### FTIR on Isolated Starches

In order to confirm the starch nature of ASC identified in ASC-2 and ASC-3 samples, we focused the analysis on ASC-4. In Figure 5K, the FTIR spectra of four isolated starches (ASC-4), three from Brinzeni I site (#442, #6707, and #3539) and one from Surein I, are shown. Spectral analysis and comparison with the vibrational profiles in Figure 5I allow us to confirm the findings and assignments done on the previous dataset on ASC-2 and ASC-3 samples. All spectra exhibit peculiar carbohydrate features, associated to −OH stretching, −CH stretching and ring vibrations in the spectral region 1200-900 cm^−1^, and relative intensity and positions of the bands can vary sample from samples, as already highlighted. Furthermore, as observed in other ASC-2 and ASC-3 (e.g. Figure 5I-e and 5I-f, black and red lines), and the ASC-4 starch from BZ#442 (Figure 5K-d black line) underwent a partial mineralization process because the signals from CaCO3 are also evident. Since the presence of the extinction cross (Figure 5 L-a) undoubtedly confirms the starch nature of ASC-4 remains and the optical images exclude a severe cross-contamination from soil particulate, it is possible to conclude that the mineral/carbonate features are peculiar of ancient starches and indicative of the diagenetic process they underwent.

In Figure 5L are shown the optical images of a starch isolated from BZ#2965 (a) and one from BZ# 6707 (b) along with their cross polarization images obtained on the glass slide and below are presented the images of the same ASC-4 after being transferred onto the ZnSe optical window.

### FTIR on modern starches

It has to be highlighted that the spectral variability in the 1200-900 cm^−1^ spectral region, particularly evident for the Brizeni samples #442, #6707, and #3539 in Figure 5K, could be also due to a different origin of the isolated starches. Since it is likely that modern plants changed over a period of 40.000 years ^1^, the former hypothesis cannot be verified by a direct comparison of what was retrieved from the pestles and GSTs with non-existent standards. This is even more true if we also consider the chemical modifications due to the aging undergone by the ASCs.

Nevertheless, some similarities in the ASCs spectra could be observed and used to classify them. To this aim, we acquired data on modern extracted starches, MSR-1 and MSR-2, and used them to build a model capable of categorizing the ASCs with respect to their modern counterparts. The database contains 137 spectra of starches belonging to eleven different plant species representing both under (USO) and above (ASO) storage organs from boreal and tropical environments: rice (*Oryza sativa*), water chestnut (*Trapa natans*), broadleaf cattail (*Typha latifolia*), narrowleaf cattail (*Typha angustifolia*), nutgrass (*Cyperus rotundus*), kudzu (*Pueraria* sp.), horse chestnut (*Aesculus hippocastanum*), rapeseed (*Brassica rapa* var. *sylvestris*) from boreal biotopes, while sweet potato (*Ipomoea batatas*), manioc (*Manihot esculenta*), and yam (*Dioscorea* sp.) represent the tropical species.

The model uses a principal component analysis (PCA) to identify the spectral features of maximum variance within the dataset and a kNN (k-nearest neighbors) algorithm to classify the unknown data in the PC space. A more detailed description and the performance indicators of the method evaluated by using stratified cross validation are presented in the supplementary materials.

In Figure 6A is presented the PC1 and PC4 score-plot of the PCA space used to generate the chemometric model; these two components have been selected since they grant the best separation among the classes. In panel 6B are shown the spectral loadings of PC1 and PC4. The loadings represent the ensemble of the spectral features that account the largest variance for each component. PC1 main peaks are those of carbohydrates: 1168 cm^−1^ from ring “breathing” vibration, 1065 cm^−1^ due to in plane bending of C-OH bond and 1030 cm^−1^ assigned to the C-OH stretching ^54^. PC4, instead, presents the most intense peaks at 1730 cm^−1^ and 1704 cm^−1^, assigned to the C=O stretching from oxidation products and aldehyde group, 1613 cm^−1^ and 1567 cm^−1^ from the stretching of the vinyl bonds, and a peak at 1423 cm^−1^ from carbonates.

**Fig. 6.**
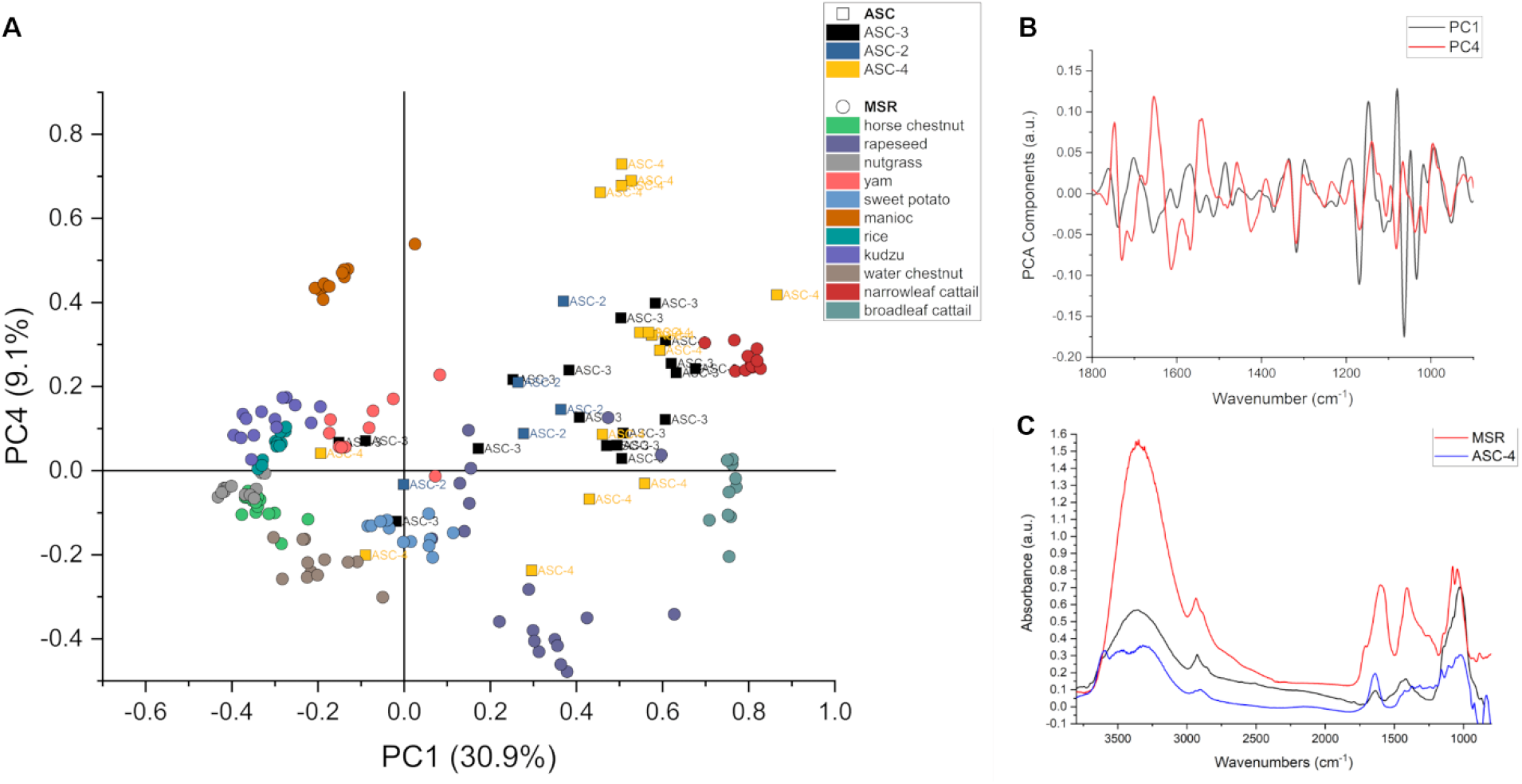
Chemometric model developed for ASCs classification. **(A)** Scatterplot of the MSR (spheres) and ASC (squares) in the PC1-PC4 space. **(B)** Spectral components representing PC1, in black, and PC4, in red, in the spectra range 1800-900 cm^−1^. **(C)** Comparison of a spectrum of an ASC-3, in red, ASC-4 in green, and one MSR (*Manihot esculenta*) in blue. The highest clustering is within the USOs (roots, tubers, and rhizomes), compatible with the presence of geophytes in the Pontic Steppe.

In Fig. 6A are also included the tested ASCs (4 ASC-2, 19 ASC-3 and the 13 ASC-4), not used for the model, in order to make possible the visualization of the relationship of the spectra of the ASCs in respect to those of the MSRs. It can be seen that most of the ASCs are in a PC1-PC4 positive quadrant, with some of them a little farer from the references, that are the spectra of the ASCs more affected by the aging and mineralization processes. Those better preserved are closer to those of the MSRs. This can be better appreciated in Fig. 5C, where we present a comparison between ASC-3, ASC-4, and MSR representative spectra. It can be noticed that both ASCs lost most of the sharp peaks in the 1200-900 cm^−1^ range, as a result of the degradation and loss of ordered structures. Another signal of aging is the loss of the intensity of the OH band, that in most of the ASCs’ spectra is lower than in the MSR, as a consequence of a withering of the plant material.

By using the built model, we identified 4 ASC-2, 19 ASC-3 and 13 ASC-4, by giving a degree of similarity to the species that these starches could belong to. Results are reported in the table in supplementary materials. From the data in Table T1 it can be seen that the majority of the starches belong to rhizomes and tubers (e.g. USOs), and only one is identified as rice (i.e., outlier in the model). It has to be noted that the partially mineralized starches from BZ#442 were also classified with a percentage of >80% as originating from a rhizome.

## Discussion

The role played by plants in the human diet is widely accepted since early times as demonstrated by the outstanding record of macrobotanical remains in the Acheulian site of Gesher-Benot Ya’aqov in Israel ^55^. However, direct data of their intentional processing for consumption is scant and mostly circumstantial ^5–7^. Late Pleistocene evidence consisting of a few grains of starch was reported from dental calculus ^1,56–58^ although their interpretation as having a food origin and exposed to thermal treatment was contended ^26^. The presence of entrapped granules in the calculus might be connected with alternative sourcing, namely the widespread ethnographic use of stomach chyme, a practice reported by Darwin (1839) during his “The Voyage on the Beagle” while observing gauchos from the Argentinian Pampa ^59^ and as recently noted for the Eskimos ^60^. Therefore, an alternative explanation for the presence of plant remains (starches and phytoliths) in dental calculus may need to be considered: it is not obviously always related to intentional starch processing.

The occurrence of use-related starchy residues on Early Upper Palaeolithic stone tools is poorly investigated and only a few starch grains have been reported on stone tools from Crimea ^12^. The wear-traces and the starch grains retrieved on ground stones dating back to the Gravettian ^13,14,61^, consistently support intentional plant processing during the Late Pleistocene and make obvious the role played by vegetable foods as part of *HS’* nutritional strategy. Indeed, our study focuses on macro-tools (not obviously modified stone tools) such as grinding stones and pestles retrieved in Aurignacian layers of two key sites for the Pontic Steppe colonization: Surein I on the Crimean Peninsula, and Brinzeni I, one of the sites that inspires the identification of the “Prut river culture” in Moldova, and referred to the earliest presence of modern humans in the area ^62^. By investigating use-related starch granules extracted from Aurignacian ground stones the present contribution has (i) established a new investigative procedure that combines morphologic and chemical analyses; (ii) highlighted proxies to identify genuine ancient starch grains, and (iii) shed light on the origins of starchy food processing in Eurasia.

We applied a strict control since starch extraction (peel-off cleaning and sonication of specific used areas) and along the whole investigative pipeline (coupling MO, SEM, FTIR). As already demonstrated by Hart ^45^ putative contamination from soil and mismanagement of stone tools may be limited to the external surface, not affecting the reliability of granules and other U-RBRs entrapped at the bottom of crevices, holes, and other uneven portions of the coarse grinding stone surfaces. We recursively optimized sample preparation and measures to obtain more reliable samples and data. As presenting the results, several different pieces of evidence and findings were piled up to support the identification of genuine ancient starchy residues from Aurignacian ground stones, thus providing key clues for the dietary breadth of modern humans during their early colonization of Eurasia, around 40.000 years ago ^28^.

One such line of evidence is the clear damage to the starch grains recovered in the samples from Brinzeni I and Surein I, visible using both optical and scanning electron microscopy. Damages to starch grains can be a result of biodegradation due to acidic or enzymatic actions due to fungi, bacteria, and enzymes ^46,47^, and/or due to mechanical processes ^43^. In our samples we observed many broken starch grains, and on many we noticed pits, cavities, the presence of conical craters, the solubilization of the internal portion of the grains, and grains exposing their lamellar structures (Figure 4 E-H) putatively interpreted as the result of enzymatic attack ^47^. Damages such as radial fibrillar fractures on the surface may be attributed to crushing, perhaps as a result of mechanical forces employed when grinding plants (Figure 4 A-O). Moreover, the co-occurrence of starch grains together with other plant elements-phytoliths, raphides, fibers, and parenchyma-not only indicates these ground stones were used to process plant matter (Figure 4 P-X), but also lessens the probability that the presence of starch grains are the result of modern contamination.

By adding the FTIR imaging it was possible to identify and characterize ASCs in complex matrices, even while accounting for the interference of mineral contaminants, and we added another piece to the puzzle allowing for the filtering of the false positives. Moreover, the SR-FTIR data on ASC-4 confirmed the identification of the ASC2 and ASC3 and strengthened the attribution to genuine starch material. The possibility to collect data from a single starch grain allowed to highlight subtle differences between each inspected particle to reveal traces of biomineralization in some of them, supported by the presence of strong carbonate peaks and clay bands. Moreover, FTIR data are in agreement of the SEM findings: some ASC-3 spectra show signals of calcium oxalate, in our data a shoulder at 1322 cm^−1^, ascribable to raphides, whereas, in other spectra, signals from lignin (1511 cm^−1^ band) and other parenchyma components are present, such as ferulic and p-coumaric acids, compound phenols which are present in plant cell walls, and which become bioavailable through gut microbiome fermentation in the small intestine ^63^. Ferulic acid owes its name to muskroot or giant fennel, *Ferula moschata*, and is common in species such as ginseng roots (Apiaceae family) and horse gram pulses (Fabaceae family).

By applying multivariate spectral analysis on modern starches spectra (reference collection built on the potential edibility and availability during the investigated period) it was also possible to propose the attribution of the ASCs we detected to genuine ancient starch grains, mainly originating from USOs, although their taxonomic attribution is still difficult to achieve ^12^. By the PCA representation in Fig. 6A and the spectra shown in figures 5I, 5K and 6C, it is clear that most of the ASC spectra collected present only a similarity with those of MSR, used to build the model, e.g. spectrum c in figure 5I and MSR in figure 6C.

This is due to all the chemical and mechanical actions that these particles were subjected to during the passing of time, detectable in the FTIR spectra as extra peaks, band shifts and bands’ component intensity variations. Nevertheless, it is exactly this not-so-perfect match with the MSR that conveys the final evidence to support the ancient origin of the retrieved particles.

Thus, the outcomes of our experiment contributed new proxies in building solid and measurable evidence for the detection and characterization of dietary carbohydrates (starch granules) processed by *HS* as soon as they arrived in the Pontic Steppe around 40.000 years ago BP. The comparison of the spectra extracted from modern starchy plants with those obtained from the ground stones use-related starch grains confirm the presence of USOs ancient starches. USOs are the structures with the purpose to store carbohydrates (starch grains) as well as oligosaccharides like inulin, and cell wall polysaccharides (Figure 4, e.g. parenchyma and fiber, Figure S1). Ancient starch grains were not the only plant residues adhering to the ground stones; we also documented other biogenic compounds such as fibers (Figures 4 and S1). Dietary fiber is important for the nutritional value of plant-based food because after undergoing fermentation by the gut microbiome, will supply valuable nutrients notably short chain fatty acids, precursors of essential biomolecules ^64^. Furthermore, several twisted fibers were recognized on the ground stones from both sites (SOM), suggesting that plant processing covered a broader range of activities, possibly even the transformation of fiber into cordage ^11^ or weaved into baskets ^65^. A further occurrence associated with Surein I starch grains is represented by raphides (calcium oxalate crystals), which have been reported as the result of processing of USOs’ at Buran Kaya III (Crimea, layer C, associated with a *HS* burial, ^12,42^).

With everything said, we believe that our multidimensional and contextual evidence strongly supports the hypothesis that the intentional processing of plant foods played a crucial role in the dietary habits of *HS*. Geophytes are predictable, usually clump to form significant amounts of biomass, and many of them are perennial, with above surface parts high enough to be visible even under snow (e.g. *Typha* sp., *Phragmites* sp., *Arundo donax*) hence, accessible during long winters ^24^. They were part of a foraging strategy that made them energetic and nutritious food, far less risky than hunting large fatty herbivores and geophytes might have become a comfort food for *HS* under the challenging climatic constraints of the northern latitudes during Late MIS 3. Pounding a plant’s storage organs disrupts the starch granules, clearly indicated by both the wear-traces and the broken grains extracted from the ground stones herein presented. Although not directly displayed by the data gathered in the presented analysis, we can speculate that after the reduction of USOs into a coarse raw flour, further boiling of this powder into a soup would be performed, as wet-cooking leads to the disruption of the degree of crystallinity of the grains and also releases insulin. Just a thermal treatment at around 90 ºC increases the susceptibility for attack by alpha-amylase in the mouth ^66^, greatly enhancing starch bio accessibility and making it a highly nutritious food. Therefore, the mechanical and thermal treatment of starchy plants are crucial steps to make dietary carbohydrates ultimately bioavailable as glucose in the bloodstream. Starchy food is calorie-rich and nutritious, hence it might have been key to maintaining homeostasis and playing a vital role in supplying the energy leading to *HS’* evolutionary success when colonizing the northern latitudes of the Pontic Steppe Belt.

By moving towards a more precisely grounded recognition and attribution of those features characterizing starches, our research made it possible to recognize authentically old use-related starch granules. Our acquired solid data can be nicely framed into a new foraging model accounting for a stricter interplay of human-land exploitation which includes intentional processing of starchy plants, a practice that early modern humans carried along while venturing Out of Africa, highly enhancing their capacity to cope with the changing subsistence conditions. This is even more true when *HS* started moving north into a totally new territory, already inhabited by different human species and undergoing dramatic climatic downturns during the late MIS 3.

## Materials and Methods

### Materials and depository

The ground stones belong to museum collections: Surein I is curated at Peter The Great Museum of Anthropology and Ethnography in St. Petersburg (MAE-RAS) and the Brinzeni I cave stone tools are curated at the National Museum of History of Moldova.

As soon as we began carrying out the first experiments, in 2017, we acknowledged a major issue concerning the authenticity of the data. We worked on previously-washed stone tools (using water from the river located near the sites), diversely processed after their retrieval from the field, and then curated in national museums. Since G. Bonch-Osmolovski’s excavation, which he magisterially documented using cutting-edge techniques, the stone from Surein I (Crimea) was very carefully re-staged at the MAE-RAS Museum, and to date is the best preserved Aurignacian context that suggests that the space was intentionally organized. During very careful excavation carried by N. Chetraru (1963-68), at Brinzeni I cave the percussive tools were recognised as ground stones and recorded in a XY coordinates system. After the excavation, the pebbles were curated in wooden boxes and no further attention (i.e. handling) was devoted to them, to the point that they are hardly mentioned in publications.

### Scanning Electron Microscopy

Non-metallized samples were investigated at IOM-CNR with Zeiss Supra 40 high resolution Field Emission Gun (FEG) Scanning Electron Microscope (SEM) with Gemini column. The decision to not metallize the samples is due to the fact that the starches in this way might be further measured with FTIR technique or other analytical techniques. The acquisition of the images was performed at low acceleration voltage (2 kV) due to the organic/biologic nature of the samples. This approach partially avoids both the charging of the surface, leading to a good quality of the final image, and the damaging of the sample itself.

### Sample preparation for Infrared analyses on the GSTs

Gentle brushing was applied, using a new clean toothbrush each time, to remove the dust that had accumulated on the objects, either stored on shelves or in boxes. In order to reproduce the tools surface texture of the used areas at the nanometric scale and to clean the surface from possible contamination, up to three impressions were taken with a high resolution molding compound (Provil Novo light by Heraeus Kulzer), following the authors procedure ^23,67^, with the dual goal of obtaining cleared area for successive sonication and a record of molds to apply the microscopic analysis of the wear-traces. The peeling effect is assumed to remove any putative contamination and was also used as a control sample. For the SEM and FTIR analyses, molds from the 2nd peeling were used. At this stage, the ground stones can be considered clean of any recent contamination and the particles remaining should be only those pushed inside the deeper ridges and crevices of the stone. U-RBR and starches adhering to the molds were observed under SEM but vinyl polysiloxane resulted not suitable for FTIR analysis (not reflective, nor flat). Hence, we decided to retrieve the particulate by dislodging the residues by sonication of a small piece of the molds in ultrapure water. This allowed to obtain a water suspension of the particles of interest that could be then deposited on a IR transparent substrate and measured.

Ultrasonic tanks were used to extract U-RBR out of the selected areas of the ground stones (ultrasonic power 180 W, 28 kHz is used for overall clean) at room temperature. As well, the molds (or selected parts of them) were sonicated (40 kHz for precise clean) to extract starch grains (ASC-3) and the procedure is detailed elsewhere ^23,37^.

### ASCs extraction and isolation

The implements from Surein I and Brinzeni I underwent different starch extraction procedures, all with the same aim to dislodge and to loosen the adhering nanoparticles from the cleaned areas. Surein I grinding stone and two refitting ground stones from Brinzeni (#442 and # NN; #833 and #2965 notably the grinding stone and the pestle) and 2 other implements (# 177 and # NN from square 11j) were all soaked in ultrapure water and sonicated in standard sonic-tanks with an average 20-40 kHz at room temperature for 15’ minutes at the (IHMC-RAS St. Petersburg and Institute of Chemistry, Chisinau, Moldova). Compared to other published techniques (pipetting solvent, or washing uncontrolled broad areas ^14^), precise area sonication presumably extracts material which is more likely to be ancient as the cavitation effect is more intense, and therefore removes sediment and entrapped organic particles from crevices and cracks of the ground stone. The sediment from sonicated artefacts was transferred into a 50 mL plastic test tube and concentrated by centrifuging. Other samples from Brinzeni I (a second sampling of #833, #442, and new sampling #3539, #6707, and #2965) were instead soaked in carbonated water overnight. The following day, the pellet was pipetted and transferred to a small vial. A drop of ethanol was added to each vial to stabilize the samples and no staining agents were added. The samples were then sent for starch grains extraction and analysis to two separate laboratories: IHAE-FB-RAS in Vladivostok and MSH Mondes in Nanterre, Paris. The laboratory methods followed those outlined by scholars ^41,68–70^ and used by one of the authors in other studies ^23,24,71^. Prior to analysis, all the laboratory consumables were washed using Alconox or bleach. The two laboratories used two different heavy liquids to separate the starches: CsCl in IHAE-FB-RAS in Vladivostok and the sodium polytungstate in MSH Mondes in Nanterre, France. Typically, samples are mounted on slides with a water and glycerin (1:1) solution. For the purpose of this research, which integrates both optical, SEM observations and further chemo-profiling of the starch grains (i.e. FTIR), two different slides were prepared. A set of slides were mounted dry (without glycerin) to avoid adding additional elements to the FTIR spectrum and to be used for SEM scan, while the other set was mounted with glycerin for standard optical microscopy observation. The starch grains were observed under a cross-polarized microscope (Nikon Eclipse E600 Pol) and photographs and measurements taken using the software NIS-Elements (located at the Archéoscopie Platform at the MSH Mondes). The microscope used in Vladivostok is a Zeiss AXIO Scope A1 with magnification up to 800x, while in France magnification reached 600x. Photographs were taken in both cross-polarized and transmitted light (Fig. 3). A third set of archeological starches was prepared by MSH Mondes by dropping directly on ZnSe windows. The starch isolation processes were fine tuned in order to guarantee a low interference on FTIR measurements. When dried, both SPT and CsCl form a thick (a couple of microns) layer of salt that covers the whole droplet area. Although neither of the two chemicals have strong chemical features in the mid IR region, they partially hinder the identification of ASCs and enhance the scattering and dispersion effect of the smaller ones. Therefore, the protocols were adjusted with additional rinsing steps to remove the layering left behind. Moreover, we also experimented with adding several drops of the same sample on top of each other on the same ZnSe holder in order to increase the putative number of ASCs recovered, highly enhancing the chance to hit the beam on a larger number of starch grains.

### Modern starch reference collection

With the aim of building a tailored reference collection, a sequential water/ethanol extraction from the USOs and ASOs selected according to pollen and plant lists available for the Pontic steppe ^42,72,73^ were carried out as follows at DAIS, Ca’ Foscari University of Venice. 5 grams (or multiple aliquots) of each of the raw storage organs were grinded by means of a blender (marca del minipimer) and the chopped residue was soaked in 200 mL of ultrapure water (with a resistivity of 18.2 MΩcm, obtained with the Milli-Q^®^ system Millipore) for 3 hours. The mixture was then filtered off in a Millipore filtration apparatus (Merck Millipore Glass Vacuum filtration system) by using a metallic filter with pore size of 0.1 mm. The filtered water containing the starches was then centrifuged at 7000 RPM (RCF = 6147) for 10 minutes at 20 °C, the pellet obtained by removing the supernatant were washed with 100 mL of ultrapure water and the mixture was sonicated for 10 minutes before recovering the pellet by further centrifugation. The precipitated matrix was then washed with EtOH (Sigma Aldrich), followed by the sonication and centrifugation processes as described above. The pellets recovered after this extraction procedure were dried and added to the permanent reference collection. An aliquot of 1 mg was dispersed in ultrapure water and droplets of 0,5 μL were set on the ZnSe holder for SR-FTIR analyses and SEM inspection.

### Infrared measurements

Samples were measured at SISSI (Synchrotron Infrared Source for Spectroscopy and Imaging) at Elettra – Sincrotrone Trieste ^74^. The molds from the second peeling of the grinding stones and pestles were sonicated in ultrapure Milli-Q^®^ water to retrieve the starch material (ASC-3), along with any other powder material present on the mold’s surface. The water suspension so obtained was centrifuged and concentrated in a 1.5 mL Eppendorf vial. 100 mL drops of the pellet were deposited onto ZnSe optical windows and dried in a sterile laminar flow hood. The samples so prepared were measured using a Bruker Hyperion 3000 IR/VIS microscope coupled with a Bruker Vertex 70V interferometer. The microscope is equipped with a 64×64 pixel Focal Plane Array (FPA) detector capable of acquiring a full FTIR spectrum per pixel, thus generating 4096 pixels’ hyperspectral images for each measure. Given the 15x magnification of the microscope, the pixel size is 2.6 x 2.6 microns and the field of view of one image (tile) is 167×167 microns. Mosaics containing multiple tiles were acquired for each sample. The measurement parameters used for each FPA measurement were the following: 64 scans at 8 cm^−1^ spectral resolution, 5 kHz scanner speed. A total of more than 50 hyperspectral maps were collected. Once areas of interest were identified by FPA imaging. Isolated starches were measured using Infrared Synchrotron radiation (IRSR) and a Mercury Cadmium Telluride (MCT) detector. The parameters used for each MCT measurement were the following: 512 scans at 4 cm^−1^ spectral resolution, 120 kHz scanner speed, setting the apertures at the same size of the starches, from 20 to 40 microns. Data was analysed in OPUS (Bruker Optics) and Quasar (https://quasar.codes) ^75,76^. Due to contamination from surrounding material some ACS’s spectra had to be further processed by subtracting the spectra of the soil background, which was obtained by extracting an average spectrum of the adjacent pixels. By analysing several samples obtained from different molds (~50 FTIR images), it was decided that in order for it to be considered a starch-candidate, the particle and its average spectrum, should have the following characteristics:

- Be a hotspot in the chemical distribution image for both −OH stretching and CH2-CH3 carbon hydrogen stretching moieties, since for C-O-C the signal could be affected by minerals; and
- Have a roundish shape with a diameter between 10 and 40 microns, even though aggregates could not be excluded

The above described conditions have the purpose to limit the areas to be analyzed. Then, once the possible spots are identified, the extracted average spectrum has to be “starch-like”^50^.

## General

We are thankful to the Director of MAE RAS, St. Petersburg (Y.K. Chistov), the Institute of Archaeology (V. Dergachev and V. Bicbaev) and M. E. Tkachuk who allowed the sampling and sustained our research, under formal MoUs and Research Agreements, and R. Ciancio (IOM-CNR) for the access and technical support to the SEM facility at IOM-CNR, Trieste, Italy. We acknowledge Elettra Sincrotrone Trieste for providing access to synchrotron radiation facility (beamtime numbers 20170057 and 20190310). We are sincerely grateful to B. Demarchi and Hoi-Ying Holman (Lawrence Berkeley National Laboratory) for critical enhancement of the earlier draft.

## Author contributions

L.L. conceived the study collaborating with N.N.S., and together with L.V. and G.B. designed the experiments. G.B., E.B., A.M., C.L., L.V. and L.L. further developed research implementation and methodological refinement; I.P. and C.C. performed the OM starch analysis; N.C. and C.S. performed SEM analysis; GS and LL developed the Digital microscopy investigation. G.B. and L.L. designed and wrote the article with input from C.C. All authors reviewed, commented and approved the final version of the manuscript and agree to be held accountable for the content therein.

