## Supplementary material for "Direct morpho-chemical characterization of elusive plant residues from Aurignacian Pontic Steppe ground stones"

### Supplementary Materials

*Archaeological sites*

**Surein I.** On the eastern slope of the Belbek gorge (Crimea Peninsula, near Sebastopol) the large rock shelter of Surein I (43 x 15 x 10 m) overlooks the second ridge of the Crimean Mountains at 110 m asl [1]. Among the best known Palaeolithic sites of Crimea, it was first excavated by K.S. Merezhkovski (alias Merejkowski) in 1879-1880 who retrieved 1150 flint artefacts, in part still conserved at the Historical Geology Chair in St. Petersburg University, and in part at the Department of Archaeology of the Peter the Great Museum of Anthropology and Ethnography (Kunstkamera) of the Russian Academy of Sciences, St. Petersburg, Russia (hereinafter MAE-RAS), where all the rest of the lithic assemblage from Surein I is curated. Between 1926-1929, G. Bonch-Osmolovski, a famed MAE-RAS museum curator, ran excavations in Surein I and Kiik Koba applying cutting-edge techniques and his findings of Crimean Neanderthals were presented to the world in 1925 with an article in Science News (pages xi-xii). It has to be stressed that G. B-O was a pioneer and made use of astonishingly modern techniques while excavating the shelter by setting a grid (2 x 2 m) to map the findings which are all coordinated within a XY system, and taking detailed notes, orthophotos and drawing of the structures he retrieved (e.g. including the grinding stone associated to flint artifacts and faunal remains still in anatomical connection). The Surein I stratigraphy was presented during the Second (proto-INQUA) Congress held in 1932 in Leningrad (Soviet Union). Moreover, G. B-O carefully collected all the materials related to human activities including 1000 artifacts in layer 3 (the lowermost level) and applied complex statistical analysis for the interpretation of the finds (Vasil'ev, 2008: 25-27, as reported in [1]). From this lowermost layer a molar tooth attributed to *Homo sapiens* was also retrieved, duly reported by Vekilova [1,2] but, at present, unfortunately lost. Layer 3 of Surein I, where the grinding stone was retrieved, is indisputably attributed to the Aurignacian [3,4]. New AMS radiocarbon dating on bone collagen - Beta 462734 carried out by the authors (unpublished) supports this; dating on new samples is on-going).

The complex structure in which the grinding stone's active surface was embedded into the sediment, has been carefully re-staged at the MAE-RAS and protected by a glass-case. Although it is not possible to control the several steps of the curation process that occurred over nearly 100 years, this organized composition of contextual presence of artifacts (both flaked and non-flaked) and animal bones, is one of the oldest and best preserved evidence of the intentional structuring of the (habitational) space by early *Homo sapiens*.

**Brinzeni I.** The cave opens on the Racovat river, a tributary of the Prut river, in the Edinet district, north-western Moldova. This is one of the most densely occupied areas during the Upper Palaeolithic - 32 and 27 ka uncal BP (36 000 and 30 000 years ago) - west of the Carpathians. Its Miocene limestone outcrops are rich in very high quality flint, possibly one of the reasons for the successful presence of modern humans in the territory during a crucial time for the colonization of south-eastern Europe. The cave, 17 x 14 x4 (maximum height) lays at 65 m asl and its mouth is oriented towards the north.

This site is recorded as yielding one of the most relevant Early Upper Palaeolithic assemblages within the so called "Prut River Culture"[5]. Its rich lowermost level 4, interpreted as cultural layer 3 by N.A. Chetraru and I.A. Borziak who systematically excavated the settlement over several campaigns (1963-65, 1968 and 1975 the former and 1987, the latter), has been attributed as Aurignacian, therefore to an early presence of *Homo sapiens* in the area (as supported by new AMS radiocarbon dating on bone collagen - Beta 462731, Beta 462732, Beta 462733 - carried out by the authors, unpublished). The site also yields extraordinary art works, among which is worth mentioning the ivory pendant or amulet (a definition Chetraru prefers, 2006 personal communication) retrieved in square 14 3, in the central area of the cave, in cultural layer 3.

The area underwent severe climatic downturns and the periglacial/steppe conditions supported by pollen analysis match with the occurrence of horse, reindeer and bison, large herbivores most targeted prey by Brinzeni dwellers. At Brinzeni I, it should be noted that trees are represented by *Pinus sylvestris*, *Betula*, *Ulmus*, *Tilia* and *Corylus*, while grasses are dominated by Poaceae, Chenopodiaceae, Asteraceae, Laminaceae, Brassicaceae, Cichoriaceae (all families that include plants with starch-rich storage organs), and in the marshy areas the genera *Polygonum*, *Cyperaceae* and *Typhaceae* are reported [5].

All the investigated ground stones were mapped and have been washed in the river near the site. Many of the pebbles still bear evident sediment residues adhering to the surfaces. Moreover, a carbonate crust was also still adhering to the grinding stone and pestles. From these areas with the carbonate crust, we also sampled for starch analysis by carefully removing said crust using a clean scalpel. The area was then molded and further sonicated or soaked to dislodge the putative remains.

*Experimental Design*

Plant remains are perishable and elusive, and their transformation, obvious in principle yet mostly assumed, is difficult to demonstrate, calling for cutting-edge and forward-looking strategies to be extracted from the working areas of pristine tools and then visually recognized. Starch grains - adhering to ground stones - are small (1-100 µm) and to be seen require analyses at the micro- and nanoscale. As most of the informative elements in the archaeological record are invisible to the naked eye, structural and physical-chemical characterization to identify the different use-related biogenic residues (U-RBR) such as starches, is required. This analytical framework is even more crucial for the latest phase of the Pleistocene record (40-25 ky), as starch-processing tools are still scant and data are based on the morphological study of starch grains, performed with visible light microscopy (OM/VLM, [6,7]). In our study we integrate light and electronic beams (Optical, Digital (DM) and Scanning Electron Microscopes) to increase both the detection of diagnostic structural features and grain size variation, that allows the identification of the morphometrics for a larger number of starch grains, mostly in their lowest size range (<50 µm) and of fibers. Molds were scanned with both a Hirox KH-8700 (MXB-2500REZ lens and a Keyence VHX-7000 with Zoom lens up to 2500 X, using both 2D and 3D modality, and with several SEM (Jeol, Zeiss). To further refine the analysis, we applied chemoprofiling with Fourier Transform Infrared (SR-FTIR) spectromicroscopy and imaging of the dried precipitated matrices obtained by the direct sonication of the ground stone and of their molds. Then we built a dataset with ancient starches extracted under controlled laboratory conditions to be measured with SR-FTIR and to be used as a cross-check for the chemoprofiling analysis. Lastly, a spectral database of the starches from modern plants available in the Pontic steppe was assembled and used to calculate a chemometric model capable of classifying the ASCs.

*Chemometric model*

In order to validate the spectra of the particles detected in both powder deposits and isolated starches, a chemometric model was built. Spectra starches, purified from modern plants, were collected with an FTIR spectro-microscope and a single point detector, and served as training dataset. The model was developed in Quasar (https://quasar.codes). The program works by visual programming, populating an empty canvas with objects that represent the individual operations and transformations done on the data. The data were loaded, interpolated to the same point spacing, and then preprocessed: data was baseline corrected with a rubber band, the second derivative was calculated using the Savitzky Golay algorithm using a 23 points window, then data was cut between 888 and 1830 $cm^{-1}$ and vector normalized. This data was then reduced using a principal component analysis (PCA) to 12 principal components that represent the 95% of the total variance (see Figure 5 in the main text). Within this 12 dimensional PCA space, k-Nearest Neighbor (kNN) algorithm was used with 4 points and the weighing distance calculated as Euclidean distance. The performance of the obtained model was evaluated by stratified cross validation repeated 10 times. The performance indicators are the following:

- Area under the Curve (AUC): 100%
- Classification Accuracy (CA): 97,1%
- F-1 Parameter: 97,1%
- Precision: 97,3%
- Recall: 97,1%

In table T1 the results of the classification of the ASCs done by the method.

**Table T1:**

| Particle ID | Type | Identification |
|---|---|---|
| Particle 1 _BZ833_1963_layer3_s5_11i_GST | ASC-3 | Pueraria sp. |
| Particle 2 _BZ833_1963_layer3_s5_11i_GST | ASC-3 | Manihot esculenta |
| Particle 3 _BZ833_1963_layer3_s5_11i_GST | ASC-3 | Manihot esculenta |
| Particle 4 _BZ833_1963_layer3_s5_11i_GST | ASC-3 | Manihot esculenta |
| Particle 5_BZ833_1963_layer3_s5_11i_GST | ASC-3 | Manihot esculenta |
| Particle 1_BZ ND_1964_layer3_s1_11j_Pestle | ASC-3 | Manihot esculenta |
| Particle 1_BZ833_1964_layer3_s1_12g_Pestle | ASC-3 | Typha angustifolia |
| Particle 1_BZ442_1964_layer3_s1_12g_GST | ASC-2 | Typha latifolia |
| Particle 2_BZ442_1964_layer3_s1_12g_GST | ASC-2 | Typha latifolia |

| | | |
|---|---|---|
| Particle 2_BZ833_1964_layer3_s1_12g_GST | ASC-3 | Oryza sativa |
| Particle 3_BZ833_1964_layer3_s1_12g_GST | ASC-3 | Trapa natans |
| Particle 1_BZ833_1964_layer3_s4_12g_Pestle | ASC-3 | Typha angustifolia |
| Particle 1_BZ833_1964_layer3_s1_12g_Pestle | ASC-3 | Ipomoea batatas |
| Particle 2_BZ833_1964_layer3_s1_12g_Pestle | ASC-3 | Ipomoea batatas |
| Particle 3_BZ833_1964_layer3_s1_12g_Pestle | ASC-3 | Brassica rapa sylvestris |
| Particle 4_BZ833_1964_layer3_s1_12g_Pestle | ASC-3 | Typha angustifolia |
| Particle 5_BZ833_1964_layer3_s1_12g_Pestle | ASC-3 | Aesculus hippocastanum |
| Particle 2_BZ833_1964_layer3_s4_12g_Pestle | ASC-3 | Typha latifolia |
| Particle 3_BZ833_1964_layer3_s4_12g_Pestle | ASC-3 | Typha angustifolia |
| Particle 4_BZ833_1964_layer3_s4_12g_Pestle | ASC-3 | Cyperus rotundus |
| Particle 1_ext_BZ6707_1963_layer3_q16e_GST | ASC-4 | Cyperus rotundus |
| Particle 2_ext_BZ6707_1963_layer3_q16e_GST | ASC-4 | Typha angustifolia |
| Particle1_Surein_I_Mold3_GST | ASC-2 | Brassica rapa sylvestris |
| Particle2_Surein_I_Mold3_GST | ASC-2 | Cyperus rotundus |
| Particle3_Surein_I_Mold3_GST | ASC-2 | Brassica rapa sylvestris |
| Particle 1_ext_BZ442_1964_layer3_s1_12g_GST | ASC-4 | Ipomoea batatas |
| Particle 2_ext_BZ442_1964_layer3_s1_12g_GST | ASC-4 | Ipomoea batatas |

| | | |
|---|---|---|
| Particle 3_ext_BZ6707_1963_layer3_q16e_GST | ASC-4 | Brassica rapa sylvestris |
| Particle 4_ext_BZ6707_1963_layer3_q16e_GST | ASC-4 | Ipomoea batatas |
| Particle 5_ext_BZ6707_1963_layer3_q16e_GST | ASC-4 | Brassica rapa sylvestris |
| Particle 6_ext_BZ6707_1963_layer3_q16e_GST | ASC-4 | Ipomoea batatas |
| Particle 7_ext_BZ6707_1963_layer3_q16e_GST | ASC-4 | Ipomoea batatas |
| Particle 8_ext_BZ6707_1963_layer3_q16e_GST | ASC-4 | Manihot esculenta |
| Particle 1_ext_BZ3539_1963_layer3_q16e_GST | ASC-4 | Brassica rapa sylvestris |
| Particle 1_ext_BZ3539_1964_layer3_q12j_GST | ASC-4 | Manihot esculenta |
| Particle 2_ext_BZ3539_1964_layer3_q12j_GST | ASC-4 | Ipomoea batatas |

The level of confidence of the prediction is divided in three levels:

- 0-33% “Fail” represented in RED
- 33-66% “Warning” represented in YELLOW
- 66-100% “Good” represented in GREEN.

As can be seen from the indicators, this model has good performances, and the identifications matched well our hypothesis of these pestles and GST mainly being used for the transformation of USOs.

Although it has some limitations that will be addressed. First, as most of the machine learning models it recognizes only the things that are in the training set, though the identifications are uncertain and it can make errors, as can be seen when Particle 2_BZ833_1964_layer3_s1_12g_GST was recognized as rice, a plant not present in the area under investigation at that time, a historical outlier placed on purpose in the database. To be more precise the database has to be as comprehensive as possible, so we have to add many more MRSs to the database. Another issue is the aging, that can alter the spectral features and make difficult/impossible the identification. For these two reasons we plan to continue our investigations by testing more starches of different origins, as a validation, and also add artificially aged samples to the database to train it also for chemically transformed starches.

*Analysis of plant fibers*

As described in the results section, in several samples, plant residues were observed (Figures 2, 3, 4 U-X). Among the many fibers retrieved, here we present one example (Figure S1), extracted during the sonication of the small fragment of the Brinzeni I grinding stone retrieved in square 12 G (small fragment). The FTIR images obtained on it and some spectra extracted from three parts of the image.

As can be seen from panels S1 A and D, the analyzed fiber thickness varies along its length, and thus the absorbance intensity varies as can be seen in Figure S1 C: in the central part (point c) is thicker and the IR signals of OH and C-O-C are stronger. From the in Figure S1 C it is possible to see that in the central part the OH band is quite narrow, hinting that all the -OH are involved in hydrogen bonds and that the structure is ordered, thus less degraded. In its narrower parts (point b in Figure S1 C-b) the band is wider, allowing us to speculate of a higher disorder of the polysaccharide chains, moreover a peak at 1740 $cm^{-1}$ of the C=O species can be seen and assigned to a partial oxidation of the plant material. Other fibers were retrieved even on molds 1, 3 and 5 of Face A of Surein I and Brinzeni I and are shown in Fig. S1 E-F.

**Figure S1.**

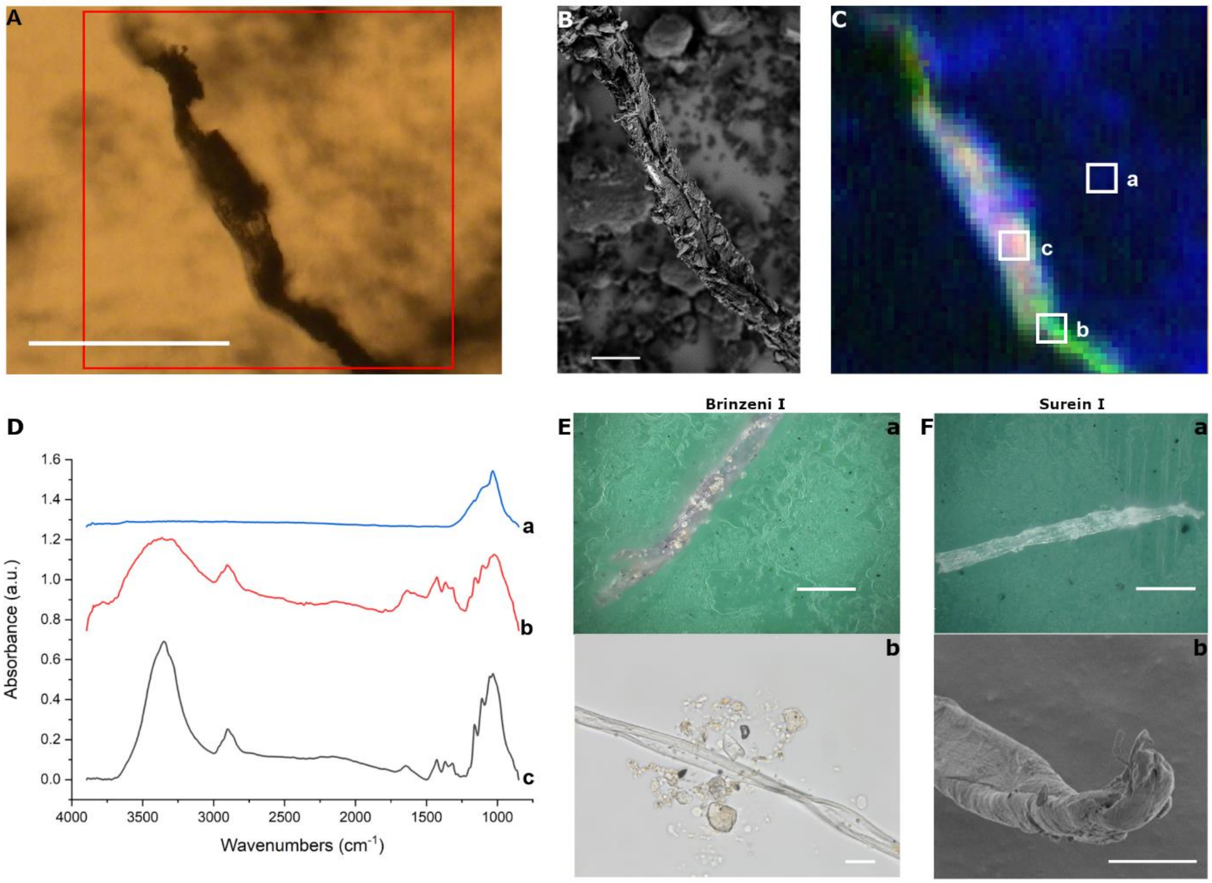


**Fig S1. (A)** Optical image of the plant fiber, the red square corresponds to the measured area. Scale Bar 100 microns. **(B)** RGB composite image of the sample: in red the integral of 3500-3100 $cm^{-1}$ band (-OH stretching), in green the integral of 3000-28000 $cm^{-1}$ band $CH_3$-$CH_2$ stretching) and in blue the integral of 1200-900 $cm^{-1}$ band (C-O-C stretching, phosphates and silicates vibrations). **(C)** Average spectra extracted from the three areas marked in panel **(C)**, **a)** mainly soil, silicates and clay, **b)** thin part of the fiber, **c)** central part of the fiber. Spectra are offset for clarity. **(D)** SEM micrograph of the fiber, scale bar 20 microns. E-F. Fibers observed associated with use-wear traces. **(E) a)** fiber on mold adhering to a polished area, as observed with a digital microscope, scale bar 50 microns, **(E) b)** fiber seen using an OM, scale bar 20 microns. **(F) a)** fiber on mold adhering to a polished area, as observed with a digital microscope, scale bar 50 microns, **(F) b)** fiber seen using SEM, 20 microns.
